# Glacial meltwater drives gene-specific diversification of metal resistance genes in High Arctic soil microbiomes

**DOI:** 10.64898/2026.06.08.730895

**Authors:** Franck J. Ouedraogo, Alexandre J. Poulain, Stéphane Aris-Brosou

## Abstract

Climate warming accelerates glacial meltwater delivery to Arctic lakes, mobilizing metals from thawing catchments and reshaping the selective landscape experienced by resident microbes. Whether these gradients leave detectable evolutionary signatures in environmental resistance genes remains unclear. We investigated four metal resistance genes (*merA, arsC, cadA*, and *chrR*) in metagenomic datasets from Lake Hazen (Nunavut, Canada), the largest High Arctic freshwater lake, sampled across a natural hydrological gradient of Control, Low-runoff, and High-runoff regimes. Using a space-for-time design, we combined population-genetic and codon-based approaches to quantify diversity and selection, including nucleotide diversity, Tajima’s *D*, nonsynonymous/synonymous diversity ratios, McDonald-Kreitman tests, site-level episodic selection, and Bayesian coalescent and structured-coalescent inference of gene-pool diversity and exchange. This multi-pronged approach allowed us to distinguish gene-specific evolutionary responses from broader demographic effects. We found marked heterogeneity among genes: *merA* showed increasing diversity and consistent evidence of adaptive evolution along the runoff gradient; *cadA* displayed the strongest adaptive signal under low runoff, with selective constraint patterns that varied across regimes; *chrR* exhibited the clearest signature of episodic positive selection, with gene-wide selection and adaptive substitutions concentrated in the high-runoff regime where chromium concentrations were greatest; while *arsC* remained largely consistent with neutral evolution across regimes. Together, these results show that climate-driven metal mobilization is associated with gene- and regime-specific diversification of microbial metal resistance gene pools in Arctic soil microbiomes. Differences in metal speciation (with arsenic and chromium occurring as redox-sensitive oxyanions, and mercury, cadmium and zinc as divalent cations) may contribute to the contrasting evolutionary trajectories observed across genes, although in-situ speciation was not assessed in this dataset. Our findings highlight environmental resistance genes as sensitive indicators of changing biogeochemical conditions in rapidly warming polar ecosystems.

## 1 Introduction

The Arctic is warming nearly four times faster than the global average, destabilizing cryospheric reservoirs that have sequestered metals for millennia (Rantanen et al., 2022; Box et al., 2019). Permafrost thaw and glacial retreat are releasing mercury, chromium, arsenic, zinc, and other metals into downstream freshwater systems (Schuster et al., 2018; Hawkings et al., 2014)., where they encounter microbial communities adapted to historically pristine conditions The biological consequences of this rapid mobilization remain poorly understood, particularly at the level of microbial gene-pool evolution. Metal resistance genes (MRGs) provide a tractable system for addressing this question because their phenotype maps directly onto a measurable environmental stressor, their molecular machinery is well-characterized, and they are often encoded on mobile genetic elements that allow rapid spread between host lineages (Bruins et al., 2000; Nies, 2003).

Four MRGs are particularly relevant to Arctic biogeochemistry: *merA* encodes mercuric reductase, which converts Hg (II) to volatile Hg (0) (Barkay et al., 2003); *arsC* encodes arsenate reductase, which reduces As (V) to As (III) for subsequent extrusion (Mukhopadhyay et al., 2002); *cadA* encodes a P-type ATPase that exports Zn (II), Cd (II), and Co (II) (Argüello et al., 2007); *chrR* encodes a chromate reductase that detoxifies Cr (VI) to less mobile Cr (III) forms (Ackerley et al., 2004). These genes encode distinct biochemical functions and differ in their genomic mobility, raising the possibility that they will respond differently to environmental metal gradients.

Lake Hazen on Ellesmere Island, Nunavut, is the largest freshwater lake by volume in the High Arctic and a sentinel site for climate-driven change (Lehnherr et al., 2018). Its watershed exhibits a natural gradient of glacial meltwater influence: the Ruggles River outflow (no glacial input), Blister Creek (low runoff), and the Abbé and Snowgoose rivers (high runoff). This gradient produces large differences in metal concentrations across regimes. Chromium increases monotonically with runoff, spanning a 60-fold range from 0.33 µg/L in the Ruggles outflow to 19.73 ± 2.24 µg/L at the Abbé and Snowgoose rivers. Zinc shows a similar monotonic pattern, ranging from below detection limits in lake surface water to 108.07 ± 12.02 µg/L in High runoff. In contrast, total mercury follows a non-monotonic pattern with maximum concentrations in the Low runoff regime (Blister Creek: 17.56 ± 14.14 ng/L) and lower but still elevated levels in High runoff (7.07 ± 1.27 ng/L vs 0.42 ng/L in the Ruggles outflow), reflecting differences in both glacial reservoir chemistry and watershed discharge volumes (St. Pierre et al., 2019). The system therefore offers a space-for-time substitution design, in which contemporary regimes serve as proxies for the trajectory expected under continued glacial retreat.

Inferring selection from metagenomic gene-pool data poses two well-known methodological challenges. First, coalescent estimators of effective population size (*N*_*e*_) assume neutrality; when selection is present, *N*_*e*_ estimates are biased downward, complicating their interpretation (Charlesworth, 2009). Second, horizontal gene transfer of MRGs via plasmids and transposons violates the vertical inheritance assumption of single-tree coalescent models (Soucy et al., 2015). We addressed both challenges by combining five complementary analyses, two of which (non-synonymous to synonymous diversity ratio *π*_N_/*π*_S_ and the McDonald-Kreitman test) do not depend on *N*_*e*_ estimation, with three codon and tree-based approaches MEME (Murrell et al., 2012), BUSTED-E (Selberg et al., 2025), BEAST2 (Bouckaert et al., 2019) / MASCOT (Müller et al., 2018).

We tested three hypotheses: (H1) nucleotide diversity (*π*) of metal resistance genes increases along the gradient of glacial meltwater input, consistent with greater metal mobilization driving the diversification of resistant variants; (H2) selective pressure on MRGs varies with environmental metal exposure, with genes responding to enriched metals showing evidence of selection that scales with concentration; and (H3) independent population-genetic methods provide convergent evidence of selection, allowing biological interpretation despite the methodological limitations of any single approach applied to metagenomic data.

## 2 Materials and Methods

### 2.1 Site and metagenomic data

Sequencing data were obtained from two published Lake Hazen surveys deposited in NCBI BioProjects PRJNA556841 (Colby et al., 2020). We restricted analyses to soil samples to focus on terrestrial microbial responses and to maintain internal comparability across the eight soil sites: CSED, CSOIL (Control regime); LR1, LR2, LRSOIL (Low runoff regime; Blister Creek catchment); and HR1, HR2, HRSOIL (High runoff regime; Abbé and Snowgoose River catchments). Reference protein sequences for *merA, arsC, cadA*, and *chrR* were retrieved from NCBI and UniProt.

### 2.2 Sequence recovery and quality control

Raw paired-end reads were quality-trimmed with Trimmomatic v0.39 (minimum quality 20, minimum length 50 bp). Putative MRG-containing reads were identified by translated BLAST against the reference protein set using DIAMOND v2.1.7 (E-value < 10^−5^, percent identity ≥ 60, query coverage ≥ 50%; Buchfink et al., 2015). We further filtered candidate sequences by enforcing the presence of conserved catalytic motifs specific to each gene family (CXXC and CPX in *cadA* (Argüello et al., 2007), the *merA* active-site Cys residues (Fox & Walsh 1983), the *arsC* redox-active CX5R (Mukhopadhyay et al., 2002), and the *chrR* FMN-binding motif), removing reads with truncated or atypical motif architecture. Sequences were translated into protein, dereplicated at 100% identity, and aligned in codon mode with MAFFT v7.526 (Katoh and Standley, 2013). After codon-aware filtering, the final soil-only datasets comprised 242, 1,349, 74, and 1,121 unique haplotypes for *merA, arsC, cadA*, and *chrR*, respectively.

### 2.3 Phylogenetic inference and diversity statistics

Maximum-likelihood phylogenies were inferred from each codon alignment with IQ-TREE v2.3.6 under the best-fit substitution model selected by ModelFinder (BIC criterion) and 1,000 ultrafast bootstrap replicates (Minh et al., 2020). Nucleotide diversity (*π*) was calculated using the Nei pairwise method (Nei, 1987) separately for each gene-regime combination on soil-only haplotypes, with 95% confidence intervals derived from 500 bootstrap resamples. Tajima’s *D* was computed using the same haplotype sets, with significance assessed against the empirical distribution under the standard neutral model (Tajima, 1989). Faith’s phylogenetic diversity was computed as the total branch length of the gene-regime subtree.

### 2.4 *N*_*e*_-independent selection tests

To address the circularity that arises when *N*_*e*_ is estimated from data under selection (Charlesworth, 2009), we applied two analyses whose statistics do not depend on *N*_*e*_. First, the ratio of nucleotide diversity at non-synonymous to synonymous sites (*π*_N_/*π*_S_) was computed using the Nei-Gojobori method (Nei and Gojobori, 1986), with 95% confidence intervals from 50 bootstrap resamples. Second, the McDonald-Kreitman (MK) test (McDonald and Kreitman, 1991) was applied per gene and regime using the best NCBI BLAST hit (*E*-value < 10^-20^) as outgroup (*Cupriavidus metallidurans* plasmid for *merA, Bacillus subtilis* for *cadA, Escherichia coli* for *arsC* and *chrR*). The proportion of adaptive substitutions *α = 1 −* (*P*_*N*_*/P*_*S*_)*/*(*D*_*N*_*/D*_*S*_) was computed following Smith and Eyre-Walker (2002), with significance assessed by a *χ*^*2*^ test with Yates correction.

### 2.5 Codon-based selection inference

Episodic site-level positive selection was inferred with the Mixed-Effects Model of Evolution (MEME) (Murrell et al., 2012) implemented in HyPhy v2.5.49 (Kosakovsky Pond et al., 2020), with empirical Bayes posterior probability used to identify codons with significant evidence for selection at *p* < 0.1. To complement MEME with a gene-wide test robust to alignment errors at high-*ω* sites, we applied the recently developed BUSTED-E (Branch-Site Unrestricted Statistical Test for Episodic Diversification with Error-sink; Selberg et al., 2025), which adds an error-absorbing rate class to the BUSTED model.

### 2.6 Bayesian coalescent inference of gene-pool diversity

We estimated the population mutation parameter *θ = 2N*_*e*_*μ* using BEAST (Bouckaert et al., 2019), v2.7.6 with the Coalescent Constant Population model (Kingman 1982 et al.,) reported throughout this study as a gene-pool diversity index rather than as an organismal demographic parameter. Inputs were per-gene, per-regime codon alignments. We ran three independent MCMC chains of 50 million iterations each, with 10% burn-in. Convergence was confirmed in Tracer (effective sample size > 200 for all parameters) and inter-chain consistency by coefficient of variation < 5%. Inter-regime gene flow was estimated using the structured coalescent model MASCOT (Müller et al., 2018), with all four genes analyzed using the same MCMC strategy.

### 2.7 Statistical analysis of metal-diversity relationships

Associations between *π* and metal concentrations were tested via Spearman rank correlation between gene-specific *π* values and the concentration of the target metal (*merA*-Hg, *arsC*-As, *cadA*-*Zn, chrR*-Cr) across the three regimes (Colby et al., 2020; St. Pierre et al., 2019Bayesian probabilities *P*(*π*_X_ > *π*_Y_) were computed from the bootstrap distributions of *π* as the proportion of bootstrap replicates in which *π*_X_ exceeded *π*_Y_, providing a posterior-like measure of directional credibility in the absence of formal Bayesian modeling. PERMANOVA on weighted UniFrac distances tested the effect of regime on phylogenetic community structure (999 permutations). All custom scripts and configuration files are available at the data availability link below.

## 3 Results

### 3.1 Haplotype recovery and overall diversity patterns

After motif-based filtering and dereplication, soil samples yielded 2,786 unique metal resistance gene haplotypes: 242 for *merA*, 1,349 for *arsC*, 74 for *cadA*, and 1,121 for *chrR* (Table 1). Recovery varied substantially across regimes for each gene, reflecting both natural abundance and sequencing depth. Nucleotide diversity (*π*) ranged from 0.044 (*merA*, Control) to 0.423 (*cadA*, High) and showed three distinct response patterns across the runoff gradient: *merA* increased monotonically with runoff (Control 0.044 → Low 0.069 → High 0.089; Bayesian *P* (*π*_*H*_ *> π*_*C*_) = 0.994); *cadA* increased with a maximum in High (0.310 → 0.296 → 0.423; *P* (*π*_*H*_ *> π*_*C*_) = 0.992); *arsC* peaked in Low (0.190 → 0.237 → 0.203) consistent with an intermediate-disturbance-like pattern; *chrR* showed a striking non-monotonic response: maximum in Low (0.240) and minimum in High (0.130), despite chromium concentrations being highest in High (Figure 1). Tajima’s *D* values supported these contrasts: *merA* exhibited negative values consistent with directional selection or expansion (*D* = -1.06 to -1.68), *arsC* and *chrR* positive values in some regimes consistent with balancing selection or population contraction.

**Table 1.**
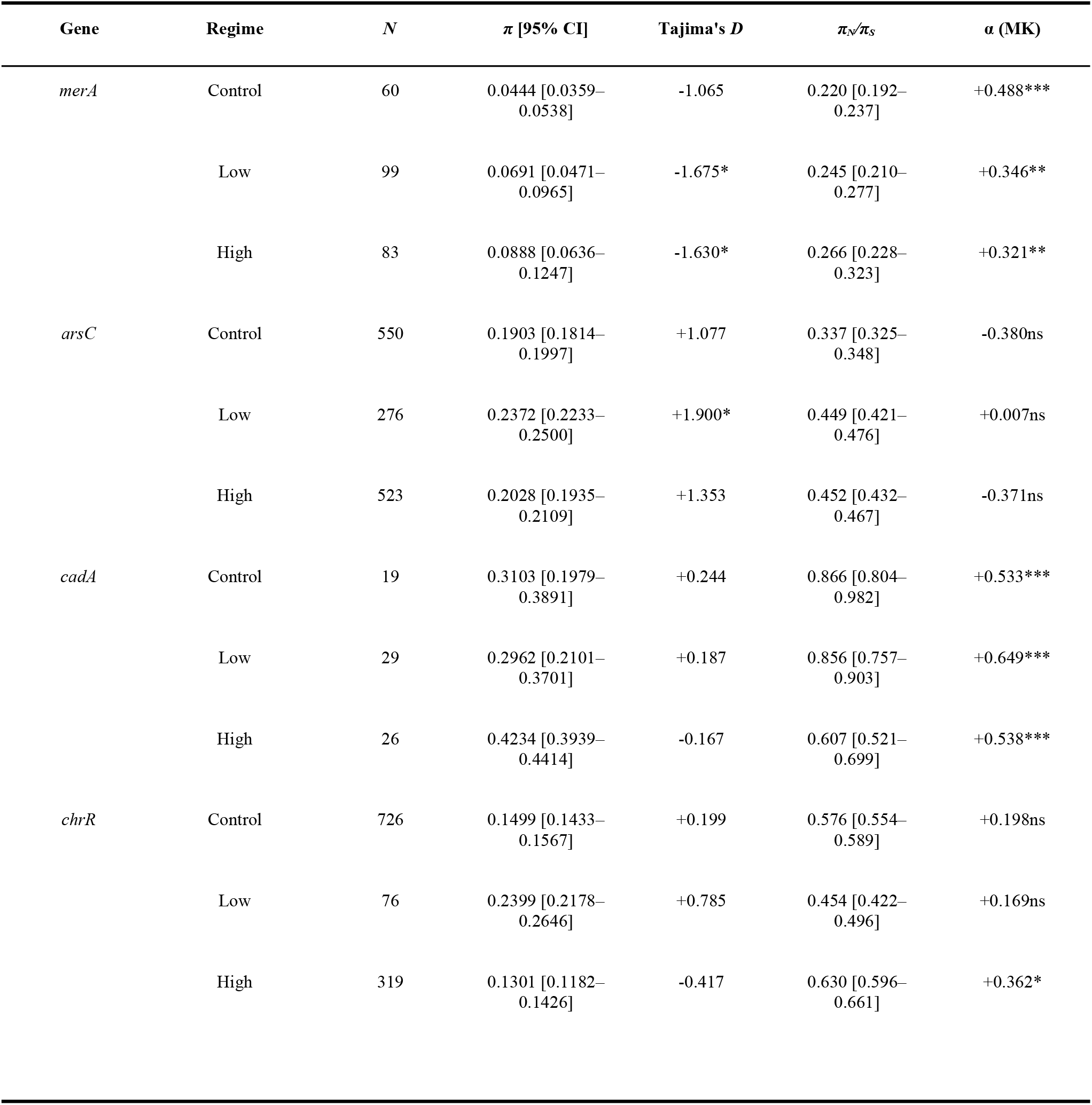
Summary statistics for the four metal resistance genes across hydrological regimes at Lake Hazen. *N* = number of unique soil haplotypes. *π* reported as mean [95% bootstrap CI]. Tajima’s *D* significance: *** *p* < 0.001, ** *p* < 0.01, * *p* < 0.05. *π*_*N*_*/π*_*S*_ = Nei-Gojobori ratio with 95% bootstrap CI. α = McDonald-Kreitman proportion of adaptive substitutions; significance from *χ*^2^ with Yates correction.

**Figure 1.**
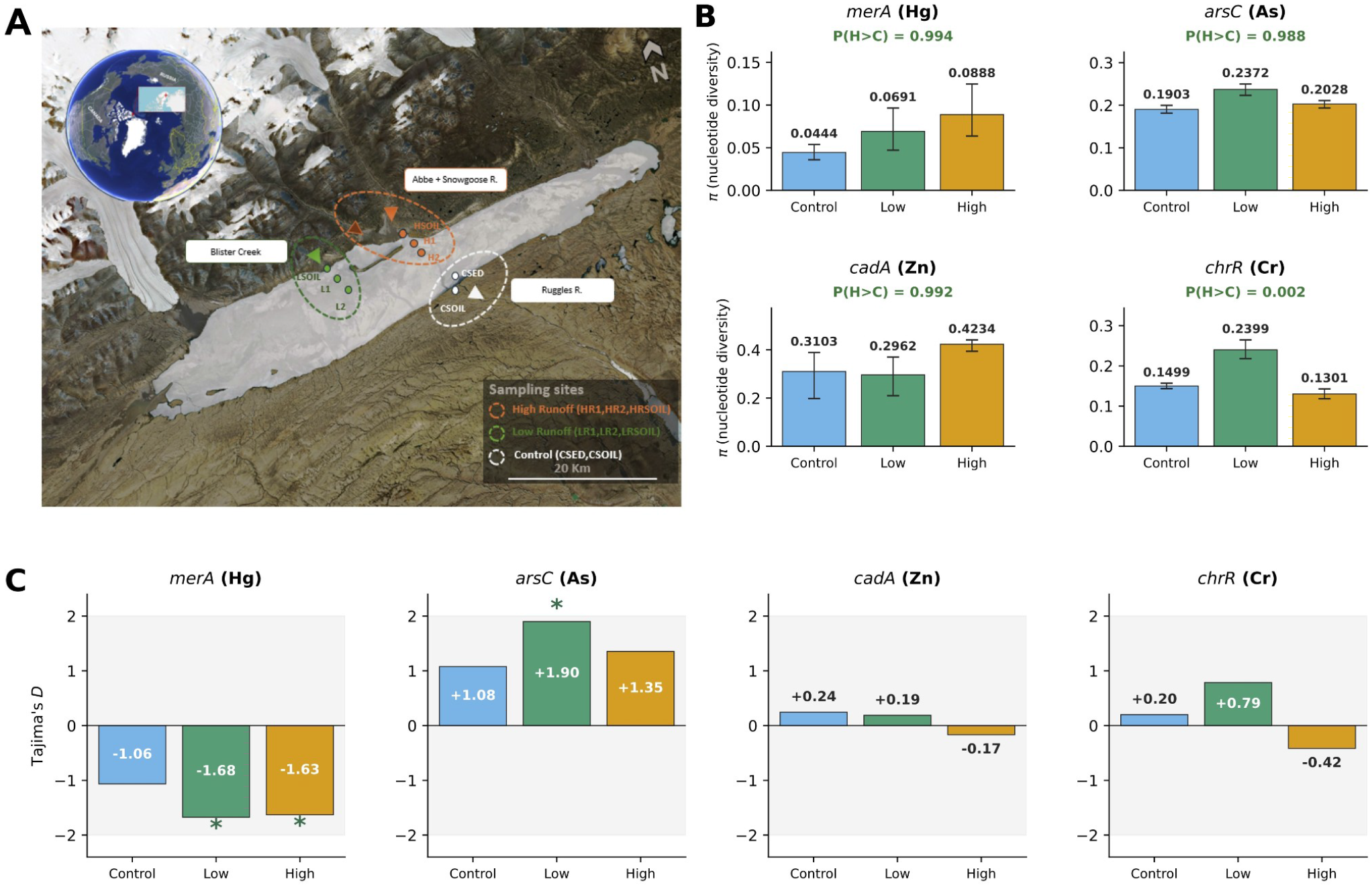
Geographic context and nucleotide diversity (*π*) of metal resistance genes at Lake Hazen.(A) Map of the Lake Hazen watershed showing the three hydrological regimes: Control (no glacial input; Ruggles River), Low (Blister Creek), and High (Abbé and Snowgoose rivers). Adapted from Colby et al. (2020). (B) Nucleotide diversity (*π*, Nei pairwise method) per gene and regime computed on soil-only haplotypes. Bars: regime mean; error bars: 95% bootstrap CI (500 resamples). Inset: Bayesian probability *P*(*π*_*H*_ *> π*_*C*_). (C) Tajima’s *D* per gene and regime. Grey band: neutral expectation (|D| < 2). Significance: *** *p* < 0.001, \*\**p* < 0.01, * *p* < 0.05, *ns = not significant*.

### 3.2 Differential selective constraint independent of Ne (*π*_*N*_*/π*_*S*_)

Nei-Gojobori *π*_N_/*π*_S_ ratios revealed gene and regime-specific patterns of selective constraint (Figure 2A; Table 1). All four genes showed ratios below 1 in all regimes, indicating dominant purifying selection across the dataset: *merA* exhibited the strongest constraint (0.220, 0.245, 0.266 across Control, Low, High), with a slight monotonic relaxation along the runoff gradient; *arsC* showed an increase from Control (0.337) to metal-exposed regimes (Low: 0.449; High: 0.452), indicating relaxed purifying selection under metal stress; *cadA* showed striking regime-specific reinforcement: *π*_*N*_/*π*_*S*_ dropped from 0.866 in Control and 0.856 in Low to 0.607 in High, a 30% reduction consistent with strong purifying selection on the zinc-binding ATPase under high zinc exposure; *chrR* showed a non-monotonic pattern with a minimum in Low (0.454) and elevated values in Control (0.576) and High (0.630).

**Figure 2.**
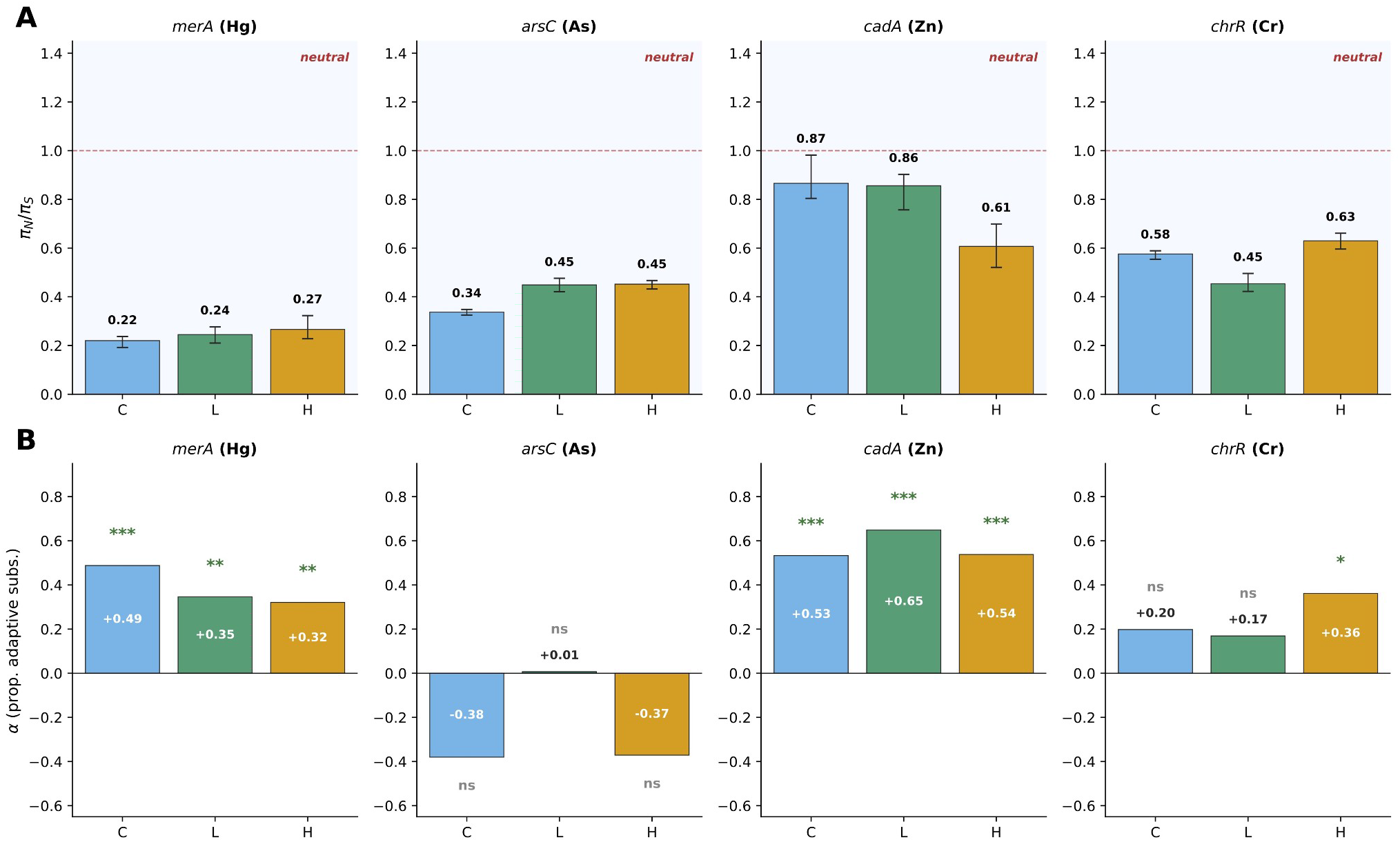
Selection inference independent of Ne estimation. (A) Ratio of non-synonymous to synonymous nucleotide diversity (*π*_N_/*π*_S_; Nei-Gojobori) per gene and regime. Error bars: 95% bootstrap CI (50 resamples). Dashed line: neutrality (*π*_*N*_*/π*_*S*_ = 1); shaded zone below 1 indicates purifying selection. (B) McDonald-Kreitman α (proportion of adaptive substitutions) per gene and regime. Significance from *χ*^2^ test with Yates correction.

### 3.3 Adaptive evolution detected by McDonald-Kreitman test

The McDonald-Kreitman test detected gene and regime-specific adaptive evolution independent of Ne estimation (Figure 2B; Table 1): *merA* showed significant excess of non-synonymous over synonymous divergence across all three regimes (*α* = +0.488, *p* < 0.001 in Control; *α* = +0.346, *p* = 0.004 in Low; *α =* +0.321, *p* = 0.007 *in High*), indicating sustained adaptive evolution along the entire gradient; *cadA* exhibited the strongest and most consistent adaptive signal, with *α* reaching its maximum in Low (+0.649, *p <* 0.001) and remaining highly significant in Control (+0.533) and High (*+0*.*538; all p < 0*.*001*); *chrR* showed the most striking regime-specific pattern: significant adaptive evolution exclusively in High (*α =* +0.362, *p* = 0.038), where chromium concentrations are highest (19.73 ± 2.24 µg/L), and no significant signal in Control (*α =* +0.198, *p* = 0.33) or Low (*α =* +0.169, *p* = 0.440). In contrast, *arsC* showed no significant departure from neutrality in any regime (|*α*| < 0.4, *p* > 0.200).

### 3.4 Codon-level and gene-wide selection (MEME and BUSTED-E)

MEME identified codon sites under episodic positive selection in all four genes (Figure 3). The proportion of sites under selection at *p < 0*.*1* was highest for *chrR* (125 of 222 codons, 56.3%) and *arsC* (86 of 162, 53.1%), and moderate for *merA* (48 of 574, 8.4%) and *cadA* (41 of 727, 5.6%). BUSTED-E confirmed gene-wide evidence of episodic positive selection for all four genes after accounting for potential alignment errors (*merA p* = 0.001, *arsC p* = 0.001, *cadA p* = 0.001, *chrR p* < 10^-7^). The error-sink class (*ω* > 100) absorbed less than 1% of site weight for all genes, indicating that the selection signal detected by MEME is not predominantly attributable to alignment artifacts.

**Figure 3.**
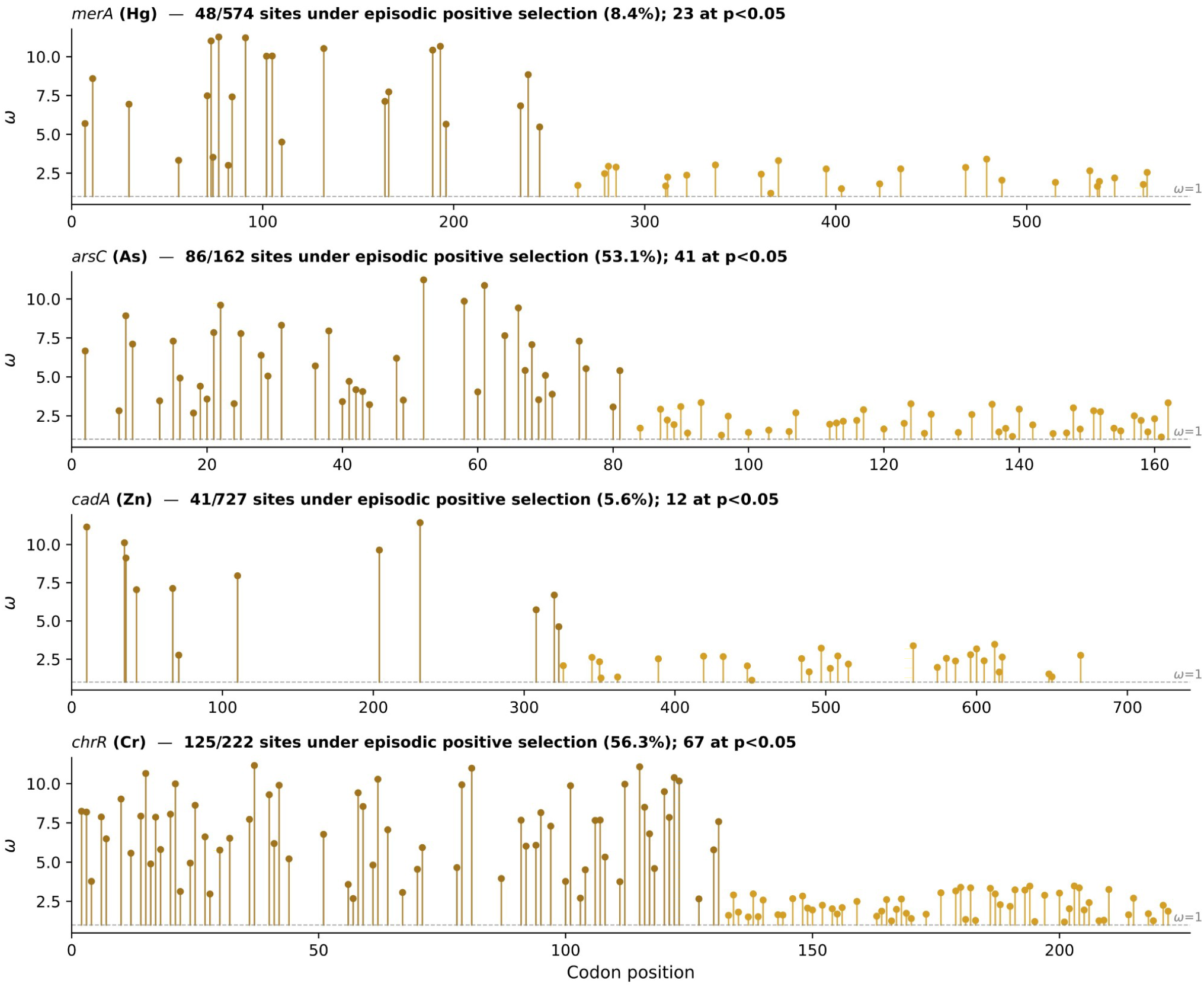
Codon positions under episodic positive selection detected by MEME (Murrell et al., 2012) per gene. Each lollipop marks a codon site with significant evidence for episodic selection at *p* < 0.1 (empirical Bayes filter). Vertical scale: per-site *ω*. Dashed line: *ω* = 1 (neutral expectation). Number and percentage of significant sites indicated for each gene.

### 3.5 Gene-pool diversity and inter-regime gene flow

Bayesian coalescent inference yielded well-converged estimates of the gene-pool diversity index *θ = 2N*_*e*_*μ* for all four genes (Figure 4A). All parameters had effective sample sizes above 1,800 and inter-chain coefficients of variation below 4%. Prior-versus-posterior comparisons confirmed the data were informative for all *θ* estimates (Figure S5), with 95% HPD width reductions of 68 to 98% relative to the Exp (1.0) prior. The *θ* rankings paralleled the *π* patterns described above: *merA* increased monotonically with runoff, *cadA* and *chrR* peaked under their target-metal regime, and *arsC* remained relatively constant across regimes. MASCOT structured coalescent analysis revealed strikingly gene-specific patterns of inter-regime gene flow (Figure 4B); *chrR* showed the lowest overall migration rates (range: 0.106 to 0.310), consistent with the strongest population structuring among the four genes; *cadA* showed the highest absolute migration rates and most symmetric pattern, consistent with its plasmid-borne mobility.

**Figure 4.**
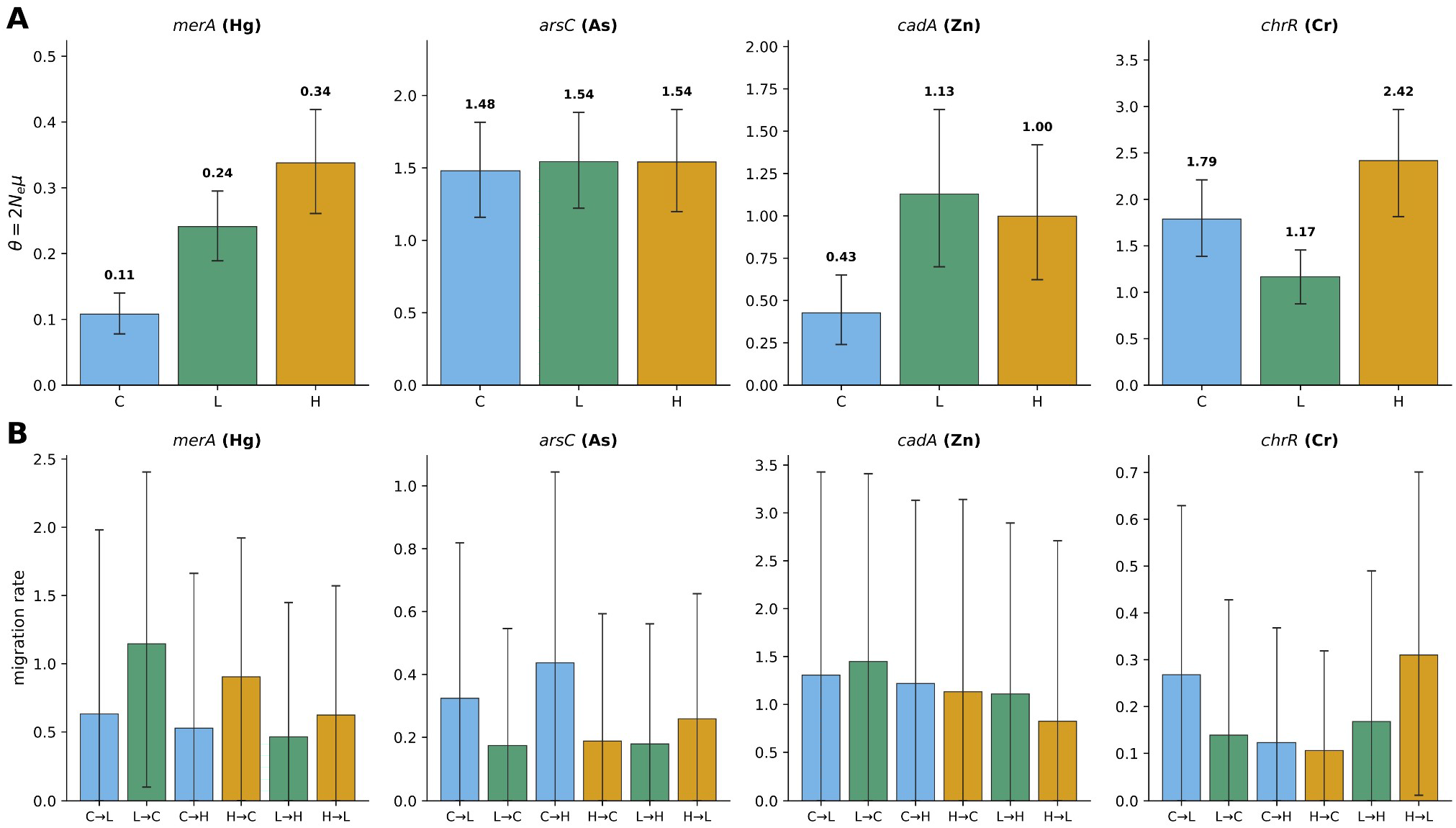
Bayesian coalescent inference of gene-pool diversity and inter-regime gene flow. (A) Posterior estimates of the gene-pool diversity index *θ = 2N*_*e*_*μ* from BEAST2 (Bouckaert et al., 2019). Bars: median; error bars: 95% highest posterior density (HPD) interval. (B) Inter-regime migration rates from MASCOT structured coalescent (Müller et al., 2018), shown for all six directional pairs. Bars: posterior median; error bars: 95% HPD.

### 3.6 Correlation between nucleotide diversity and metal concentrations

Spearman rank correlations between regime-level *π* and corresponding metal concentrations revealed gene-specific associations (Figure 5): *cadA π* and zinc showed *ρ =* +0.50, with the highest *π* in High where Zn concentrations are highest; *arsC* and As also showed *ρ =* +0.50, reflecting a partial monotonic relationship. *merA* showed *ρ = +0*.*50* with THg, but the relationship is shaped by the non-monotonic THg concentration pattern (Low > High > Control) combined with the monotonic increase of *merA* diversity along the runoff gradient: *π* tracks the integrated mercury flux rather than instantaneous concentration. *chrR* showed a negative correlation with chromium (*ρ =* -0.50), driven by its non-monotonic *π* trajectory (maximum in Low, minimum in High) despite the monotonic increase of Cr. With only three regimes the Spearman p-values are not statistically interpretable; the Bayesian probabilities *P* (*π*_*H*_ *> π*_*C*_) reported in Figure 1 provide complementary directional support.

**Figure 5.**
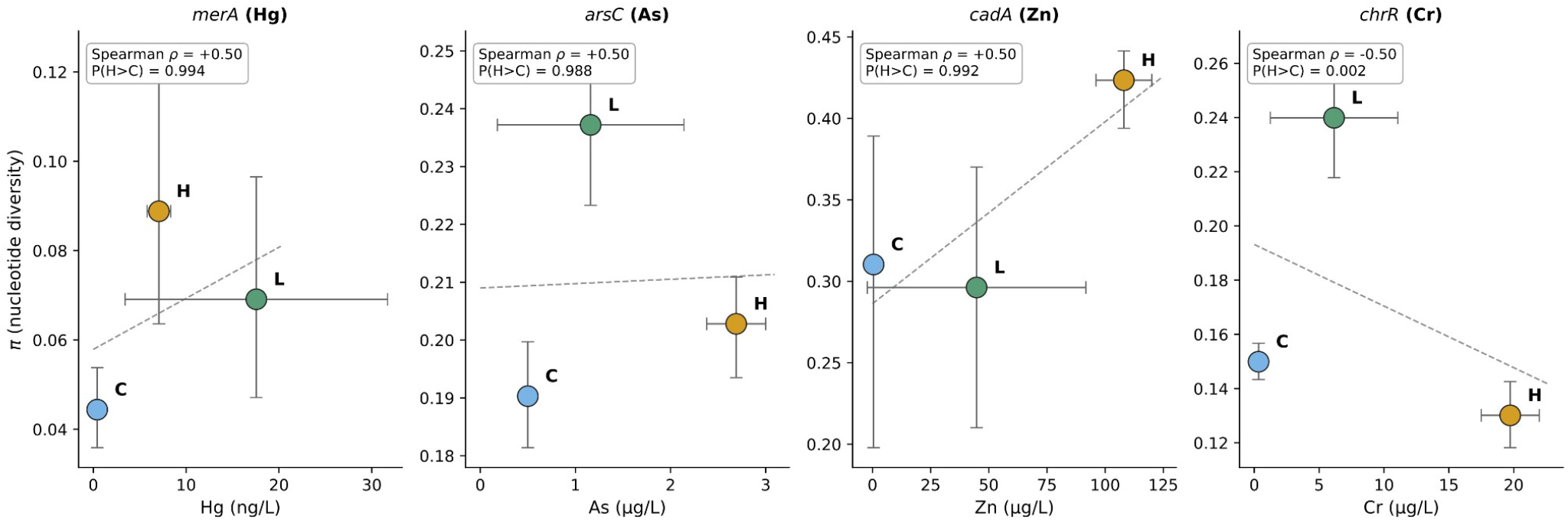
Relationship between nucleotide diversity (*π*) and target metal concentration per gene. Each point represents one regime (C = Control, L = Low, H = High). Horizontal error bars: metal SD across replicate sites; vertical error bars: *π* bootstrap 95% CI. Spearman rank correlation (*n* = 3, *p*-values not statistically interpretable with three points) and Bayesian *P*(*π*_*H*_ *> π*_*C*_) indicated in insets.

## 4 Discussion

Five complementary analyses converged on the conclusion that variable glacial meltwater inputs at Lake Hazen leave detectable, gene and regime-specific evolutionary signatures on metal resistance gene pools. Although each individual method has known limitations, signals that recur across approaches with non-overlapping assumptions can be interpreted with substantially greater credibility than those supported by any single line of evidence. This convergence allows us to interpret three contrasting evolutionary patterns across our four MRGs, which we describe in turn below.

The first pattern sustained adaptive evolution along the runoff gradient was observed for *merA*. This gene showed monotonic gene-pool diversification consistent with continuous selection pressure imposed by mercury delivery to the lake basin and with the well-documented mobility of *merA* on the Tn21 transposon family (Liebert et al., 1999). Although mercury concentrations are highest in the Low runoff regime (Blister Creek; 17.56 ± 14.14 ng/L) rather than in the High regime (Snowgoose and Abbé; 7.07 ± 1.27 ng/L), *merA* diversity tracks the runoff gradient rather than the concentration gradient. We interpret this as evidence that the standing mercury exposure of soil microbiomes is best captured by flux-based metrics, which integrate concentration over the much greater discharge volume of the Snowgoose and Abbé river systems compared to Blister Creek (St. Pierre et al., 2019). The second pattern regime-specific peaks of selection aligned with target-metal gradients were observed for *cadA* and *chrR*. For *cadA*, the simultaneous detection of strong adaptive evolution (α = +0.65 in Low; +0.54 in High) and a 30% reduction in π_N_/π_S_ in High (0.61 vs 0.87 in Control) indicates a population continually exploring variant space under selection while environmental pressure progressively narrows the functionally permissible set. For *chrR*, the locally adaptive McDonald-Kreitman signal restricted to the High regime, combined with the highest proportion of MEME-detected positive selection (56% of codons) and the lowest inter-regime gene flow of any gene, indicates a population consistent with adaptive diversification in the Abbé-Snowgoose river system but remaining structurally distinct from low-Cr regimes. The third pattern a near-absence of detectable adaptive signal was observed for *arsC*. One biogeochemical consideration is that arsenic and chromium are redox-sensitive metalloids that occur in the environment primarily as oxyanions (arsenate, arsenite, chromate), whose mobility, bioavailability, and cellular uptake depend strongly on local pH and redox potential. Their cellular accessibility therefore differs from that of divalent cations such as 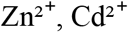, and 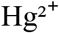, which form cationic ligand complexes and follow different transport pathways into the cell.

On this basis one might expect a priori that *arsC* and *chrR*, which both target oxyanion substrates, would behave more similarly to each other than to *merA* or *cadA*. Our data does not support that simple grouping: *chrR* shows the strongest signature of regime-specific positive selection (significant McDonald-Kreitman α only in High, lowest inter-regime gene flow, and the highest fraction of MEME-detected sites), whereas *arsC* shows no detectable adaptive signal in any regime. The contrast between these two oxyanion-targeting genes indicates that substrate speciation alone is not sufficient to predict the evolutionary trajectory of an MRG in this system, and that gene-specific factors including the magnitude of the local exposure gradient, host-cell physiology, and the operon context in which each gene is embedded must also be considered. A central limitation of our dataset is that we did not measure in-situ metal speciation or redox conditions, and our interpretation is therefore restricted to total metal pools across regimes. Resolving the relative contributions of speciation, redox chemistry, and host physiology to MRG evolution will require coupled metagenomic and geochemical speciation surveys.

Two methodological considerations frame our interpretation. First, coalescent estimators of *N*_*e*_ assume neutrality. When selection is present (as results show for all four genes), *N*_*e*_ estimates are biased downward (Charlesworth, 2009). We addressed this circularity by adding two *N*_*e*_-independent selection tests (*π*_N_/*π*_S_ and McDonald-Kreitman) and by reporting coalescent outputs as gene-pool diversity indices rather than as estimates of organismal effective population size. The convergence of *N*_*e*_-independent and codon-based tests with regime-specific coalescent patterns supports the biological interpretation that follows. Second, several MRGs reside on mobile genetic elements (Liebert et al., 1999; Silver and Phung, 1996), violating the vertical inheritance assumption of single-tree coalescent models. Two considerations mitigate this concern: the recovery of consistent gene-specific patterns across methods with different inheritance assumptions, and the conceptual framing of metagenomic gene pools as standing variation in environmental gene pools rather than as organismal lineages. The empirical alignment between MASCOT migration rates and the known mobility profile of each gene (high for *cadA*, low for *chrR*) supports this framing.

The patterns documented here have practical implications for monitoring Arctic ecosystems under accelerating climate change. The clear regime-specific selection signatures for *chrR* and *cadA* indicate that MRG gene pools can respond on timescales relevant to contemporary glacial retreat and could serve as sensitive biomarkers complementing traditional geochemical measurements. Three limitations should be acknowledged. First, the analysis covers a single watershed. Replicating the space-for-time substitution at other Arctic sites would test generality. Second, the small number of *cadA* haplotypes (*n* = 19 to 29 per regime) limits the precision of estimates for this gene, although the consistency of the MK signal across regimes provides robust evidence of selection. Third, alternative outgroup choices in the McDonald-Kreitman test could yield slightly different α estimates: the use of reciprocal best BLAST hits represents a conservative choice. Long-read metagenomics combined with genome binning would clarify the contribution of host phylogeny versus mobile element dynamics, and functional characterization of regime-specific variants under positive selection (particularly *chrR* variants in the High regime) would test the phenotypic significance of the genetic signatures inferred here. As climate-driven mobilization of previously sequestered metals accelerates across the Arctic, monitoring the evolutionary responses of MRG pools alongside chemical measurements may provide an integrative framework for assessing the biological consequences of these rapid environmental changes.

## 6 Author Contributions

FJO, AJP, and SAB designed the study. FJO performed all computational analyses and wrote the first draft. SAB supervised the evolutionary and bioinformatic analyses. AJP supervised geochemical interpretation. All authors contributed to the interpretation of results, edited the manuscript, and approved the final version.

## 8 Acknowledgments

We thank the Aris-Brosou and Poulain laboratories at the University of Ottawa for valuable discussions throughout this work. Computational resources were provided by the Digital Research Alliance of Canada (formerly Compute Canada). We thank G. A. Colby and colleagues for making the underlying metagenomic data publicly available through NCBI BioProjects PRJNA556841 and PRJNA746497.

## 9 Supplementary Material

The Supplementary Material for this article includes additional figures showing per-site nucleotide diversity, bootstrap distributions, phylogenetic diversity (Faith’s PD), the relationship between *θ* and *π*, prior-versus-posterior comparisons, and the ω distribution from MEME (Figures S1-S6), as well as supplementary tables presenting detailed per-site diversity values, Tajima’s *D* statistics, metal concentrations, PERMANOVA results, BEAST2 and MASCOT posterior estimates, MEME results, *π*_N_/*π*_S_ values, and McDonald-Kreitman test outputs (Tables S1-S10).

## Data Availability Statement

The raw metagenomic sequencing reads analyzed in this study are publicly available in the NCBI Sequence Read Archive under BioProjects PRJNA556841 (Colby et al., 2020). The trace metal and total mercury concentration data are from St. Pierre et al. (2019). All custom analysis scripts, used in this study are available in the GitHub repository https://github.com/Franckoued/mrg-evolution-lakehazen.git.

