## Supplementary material for "Glacial meltwater drives gene-specific diversification of metal resistance genes in High Arctic soil microbiomes": SI file

### 1 Supplementary Tables

**Table S1.** Nucleotide diversity ($\pi$) per gene and individual sampling site, computed by the Nei pairwise method with 500 bootstrap replicates for 95% confidence intervals. Sites grouped by hydrological regime and sample type (SED = sediment, SOIL = soil). Missing values indicate insufficient sequence recovery ($N < 5$).

| Gene | Site | Regime | Type | $N$ | $\pi$ [95% CI] |
|---|---|---|---|---|---|
| *merA* | CSOIL | C | Soil | 60 | 0.0444 [0.0359–0.0538] |
| | LSOIL | L | Soil | 99 | 0.0691 [0.0471–0.0965] |
| | H1 | H | Sed | 112 | 0.1840 [0.1634–0.2039] |
| | HSOIL | H | Soil | 83 | 0.0888 [0.0636–0.1247] |
| *arsC* | CSED | C | Sed | 10 | 0.3347 [0.2474–0.3876] |
| | CSOIL | C | Soil | 550 | 0.1903 [0.1814–0.1997] |
| | L1 | L | Sed | 56 | 0.3193 [0.2893–0.3400] |
| | L2 | L | Sed | 135 | 0.3092 [0.2957–0.3246] |
| | LSOIL | L | Soil | 276 | 0.2372 [0.2233–0.2500] |
| | H1 | H | Sed | 238 | 0.2522 [0.2421–0.2621] |
| | H2 | H | Sed | 71 | 0.2750 [0.2477–0.2989] |

| Gene | Site | Regime | Type | *N* | *π* [95% CI] |
|---|---|---|---|---|---|
| | HSOIL | H | Soil | 523 | 0.2028 [0.1935–0.2109] |
| *cadA* | CSOIL | C | Soil | 19 | 0.3103 [0.1979–0.3891] |
| | L1 | L | Sed | 6 | 0.2620 [0.1332–0.3444] |
| | L2 | L | Sed | 10 | 0.2463 [0.0946–0.3518] |
| | LSOIL | L | Soil | 29 | 0.2962 [0.2101–0.3701] |
| | H1 | H | Sed | 15 | 0.3559 [0.2608–0.4066] |
| | H2 | H | Sed | 5 | 0.2924 [0.0040–0.4132] |
| | HSOIL | H | Soil | 26 | 0.4234 [0.3939–0.4414] |
| *chrR* | CSED | C | Sed | 73 | 0.1223 [0.1053–0.1388] |
| | CSOIL | C | Soil | 726 | 0.1499 [0.1433–0.1567] |
| | L1 | L | Sed | 44 | 0.2317 [0.1879–0.2676] |
| | L2 | L | Sed | 217 | 0.1501 [0.1362–0.1658] |
| | LSOIL | L | Soil | 76 | 0.2399 [0.2178–0.2646] |
| | H1 | H | Sed | 327 | 0.1999 [0.1826–0.2163] |
| | H2 | H | Sed | 150 | 0.2397 [0.2186–0.2610] |
| | HSOIL | H | Soil | 319 | 0.1301 [0.1182–0.1426] |

**Table S2.** Tajima's *D* statistics for soil-only haplotypes with significance levels. * $p < 0.05$, ** $p < 0.01$, *** $p < 0.001$. Negative *D* values may indicate population expansion or purifying selection; positive values may indicate contraction or balancing selection.

| Gene | Regime | *N* haplotypes | Tajima's *D* | *p*-value | Significance |
|---|---|---|---|---|---|
| *merA* | Control | 60 | -1.065 | n.s. | ns |
| | Low | 99 | -1.675 | < 0.05 | * |
| | High | 83 | -1.630 | < 0.05 | * |
| *arsC* | Control | 550 | +1.077 | n.s. | ns |
| | Low | 276 | +1.900 | < 0.05 | * |
| | High | 523 | +1.353 | n.s. | ns |
| *cadA* | Control | 19 | +0.244 | n.s. | ns |
| | Low | 29 | +0.187 | n.s. | ns |
| | High | 26 | -0.167 | n.s. | ns |
| *chrR* | Control | 726 | +0.199 | n.s. | ns |
| | Low | 76 | +0.785 | n.s. | ns |
| | High | 319 | -0.417 | n.s. | ns |

**Table S3.** Metal concentrations across hydrological regimes at Lake Hazen (*mean* ± *SD*). All values from St. Pierre et al. (2019). Control = Lake Hazen surface water and Ruggles River outflow; Low = Blister Creek; High = pooled Snowgoose + Abbé Rivers. Note that THg follows a non-monotonic pattern with maximum in Low, while Cr and Zn show monotonic increases along the runoff gradient. H/C fold-change = ratio of concentration in High vs Control regime; n.d. = not determinable (Control below detection limit).

| Metal | Control | Low (Blister) | High (SG + Abbé) | Unit | H/C fold-change |
|---|---|---|---|---|---|
| THg | 0.42 | 17.56 ± 14.14 | 7.07 ± 1.27 | ng/L | 16.6× |
| Cr | 0.33 | 6.15 ± 4.92 | 19.73 ± 2.24 | µg/L | 60.2× |
| As | < 0.5 | 1.16 ± 0.98 | 2.69 ± 0.31 | µg/L | n.d. |
| Zn | < 0.5 | 44.77 ± 46.98 | 108.07 ± 12.02 | µg/L | n.d. |

**Table S4.** Faith's phylogenetic diversity (PD) per gene and regime, computed as the total branch length of the soil-only IQ-TREE phylogeny (substitutions per site).

| Gene | Regime | *N* haplotypes | Faith's PD (subs/site) |
|---|---|---|---|
| *merA* | Control | 60 | 0.7400 |
| | Low | 99 | 4.6690 |
| | High | 83 | 7.9180 |
| *arsC* | Control | 550 | 33.1910 |
| | Low | 276 | 22.0060 |
| | High | 523 | 35.5330 |
| *cadA* | Control | 19 | 5.1950 |
| | Low | 29 | 7.9410 |
| | High | 26 | 7.7010 |
| *chrR* | Control | 726 | 58.5650 |
| | Low | 76 | 13.7260 |
| | High | 319 | 38.6300 |

**Table S5.** Detailed BEAST2 posterior estimates of gene-pool diversity index ($\theta = 2N_e\mu$) per gene and regime, with 95% highest posterior density (HPD) interval and CI width.

| Gene | Regime | *N* haplotypes | *θ* median | 95% HPD lower | 95% HPD upper | CI width |
|---|---|---|---|---|---|---|
| *merA* | Control | 60 | 0.1080 | 0.0780 | 0.1400 | 0.0620 |
| | Low | 99 | 0.2410 | 0.1890 | 0.2950 | 0.1060 |
| | High | 83 | 0.3380 | 0.2610 | 0.4190 | 0.1580 |
| *arsC* | Control | 550 | 1.4800 | 1.1590 | 1.8150 | 0.6560 |
| | Low | 276 | 1.5430 | 1.2210 | 1.8830 | 0.6620 |
| | High | 523 | 1.5410 | 1.1980 | 1.9030 | 0.7050 |
| *cadA* | Control | 19 | 0.4270 | 0.2400 | 0.6510 | 0.4110 |
| | Low | 29 | 1.1290 | 0.6990 | 1.6280 | 0.9290 |
| | High | 26 | 0.9980 | 0.6230 | 1.4190 | 0.7960 |
| *chrR* | Control | 726 | 1.7880 | 1.3860 | 2.2110 | 0.8250 |
| | Low | 76 | 1.1670 | 0.8740 | 1.4550 | 0.5810 |
| | High | 319 | 2.4170 | 1.8140 | 2.9670 | 1.1530 |

**Table S6.** Detailed MASCOT structured coalescent inter-regime migration rates per gene, with 95% HPD intervals. Migration rates are in events per lineage per coalescent time unit.

| Gene | From | To | Median rate | 95% HPD lower | 95% HPD upper |
|---|---|---|---|---|---|
| *merA* | C | L | 0.632 | 0.000 | 1.981 |
| | L | C | 1.146 | 0.101 | 2.405 |
| | C | H | 0.528 | 0.000 | 1.662 |
| | H | C | 0.904 | 0.001 | 1.921 |
| | L | H | 0.464 | 0.000 | 1.449 |
| | H | L | 0.624 | 0.000 | 1.571 |
| *arsC* | C | L | 0.324 | 0.000 | 0.819 |
| | L | C | 0.174 | 0.000 | 0.546 |
| | C | H | 0.437 | 0.000 | 1.044 |
| | H | C | 0.188 | 0.000 | 0.593 |
| | L | H | 0.179 | 0.000 | 0.561 |
| | H | L | 0.259 | 0.000 | 0.657 |
| *cadA* | C | L | 1.306 | 0.000 | 3.427 |
| | L | C | 1.448 | 0.000 | 3.410 |
| | C | H | 1.219 | 0.000 | 3.132 |
| | H | C | 1.132 | 0.000 | 3.141 |
| | L | H | 1.109 | 0.000 | 2.894 |
| | H | L | 0.824 | 0.000 | 2.709 |
| *chrR* | C | L | 0.268 | 0.000 | 0.629 |

| Gene | From | To | Median rate | 95% HPD lower | 95% HPD upper |
|---|---|---|---|---|---|
| | L | C | 0.139 | 0.000 | 0.428 |
| | C | H | 0.123 | 0.000 | 0.368 |
| | H | C | 0.106 | 0.000 | 0.319 |
| | L | H | 0.168 | 0.000 | 0.490 |
| | H | L | 0.310 | 0.012 | 0.701 |

**Table S7.** PERMANOVA on weighted UniFrac distances, testing the effect of hydrological regime on phylogenetic community structure (999 permutations). $R^2$ is the proportion of variance explained by regime.

| Gene | $R^2$ | *p-value* | Permutations | Effect interpretation |
|---|---|---|---|---|
| *merA* | 0.064 | 0.001 | 999 | Weak (<0.2) |
| *arsC* | 0.232 | 0.001 | 999 | Moderate (0.2-0.5) |
| *cadA* | 0.154 | 0.001 | 999 | Weak (<0.2) |
| *chrR* | 0.718 | 0.001 | 999 | Strong (>0.5) |

**Table S8.** Detailed MEME results per gene: number and percentage of codons under episodic positive selection at $p < 0.1$ (empirical Bayes filter) and at the stricter $p < 0.05$ threshold.

| Gene | Total codons | Sites $p$<0.1 | % of total | Sites $p$<0.05 | % of total |
|---|---|---|---|---|---|
| *merA* | 574 | 48 | 8.4% | 23 | 4.0% |
| *arsC* | 162 | 86 | 53.1% | 41 | 25.3% |
| *cadA* | 727 | 41 | 5.6% | 12 | 1.7% |
| *chrR* | 222 | 125 | 56.3% | 67 | 30.2% |

**Table S9.** Nucleotide diversity at non-synonymous ($\pi_N$) and synonymous ($\pi_S$) sites computed using the Nei-Gojobori (1986) method, with 95% bootstrap confidence intervals (50 resamples) on the $\pi_N/\pi_S$ ratio. Ratios below 1 indicate purifying selection. Computations independent of $N_e$ estimation.

| Gene | Regime | $\pi_N$ | $\pi_S$ | $\pi_N/\pi_S$ | 95% CI | # codons |
|---|---|---|---|---|---|---|
| *merA* | Control | 0.0181 | 0.0825 | 0.220 | [0.192–0.237] | 478 |
| | Low | 0.0212 | 0.0867 | 0.245 | [0.210–0.277] | 478 |
| | High | 0.0288 | 0.1082 | 0.266 | [0.228–0.323] | 478 |
| *arsC* | Control | 0.0805 | 0.2386 | 0.337 | [0.325–0.348] | 142 |
| | Low | 0.1078 | 0.2403 | 0.449 | [0.421–0.476] | 142 |
| | High | 0.0939 | 0.2079 | 0.452 | [0.432–0.467] | 144 |
| *cadA* | Control | 0.1024 | 0.1182 | 0.866 | [0.804–0.982] | 462 |
| | Low | 0.1185 | 0.1384 | 0.856 | [0.757–0.903] | 389 |
| | High | 0.0802 | 0.1322 | 0.607 | [0.521–0.699] | 627 |
| *chrR* | Control | 0.0667 | 0.1158 | 0.576 | [0.554–0.589] | 194 |
| | Low | 0.0962 | 0.2117 | 0.454 | [0.422–0.496] | 190 |
| | High | 0.0530 | 0.0843 | 0.630 | [0.596–0.661] | 196 |

**Table S10.** McDonald-Kreitman test (1991) results per gene and regime. $P_N/P_S$: intra-regime polymorphisms. $D_N/D_S$: fixed differences vs outgroup. $NI = (P_N/P_S)/(D_N/D_S)$. $\alpha = 1 - NI$ (Smith and Eyre-Walker, 2002) is the proportion of adaptive substitutions. Significance from $\chi^2$ with Yates correction. Independent of $N_e$ estimation.

| Gene | Regime | $P_N$ | $P_S$ | $D_N$ | $D_S$ | $\chi^2$ | *p* | *NI* | *α* | Sig. |
|---|---|---|---|---|---|---|---|---|---|---|
| *merA* | Control | 123 | 206 | 168 | 144 | 16.85 | <0.0001 | 0.512 | +0.488 | *** |
| | Low | 221 | 319 | 159 | 150 | 8.39 | 0.0038 | 0.654 | +0.346 | ** |
| | High | 295 | 373 | 163 | 140 | 7.38 | 0.0066 | 0.679 | +0.321 | ** |
| *arsC* | Control | 130 | 140 | 37 | 55 | 1.43 | 0.2314 | 1.380 | -0.380 | ns |
| | Low | 126 | 139 | 42 | 46 | 0.00 | 1.0000 | 0.993 | +0.007 | ns |
| | High | 125 | 138 | 37 | 56 | 1.36 | 0.2429 | 1.371 | -0.371 | ns |
| *cadA* | Control | 289 | 274 | 278 | 123 | 30.57 | <0.0001 | 0.467 | +0.533 | *** |
| | Low | 316 | 323 | 251 | 90 | 52.22 | <0.0001 | 0.351 | +0.649 | *** |
| | High | 394 | 473 | 330 | 183 | 45.33 | <0.0001 | 0.462 | +0.538 | *** |
| *chrR* | Control | 167 | 176 | 71 | 60 | 0.94 | 0.3319 | 0.802 | +0.198 | ns |
| | Low | 153 | 153 | 71 | 59 | 0.60 | 0.4370 | 0.831 | +0.169 | ns |
| | High | 138 | 141 | 89 | 58 | 4.32 | 0.0378 | 0.638 | +0.362 | * |

### 2 Supplementary Figures

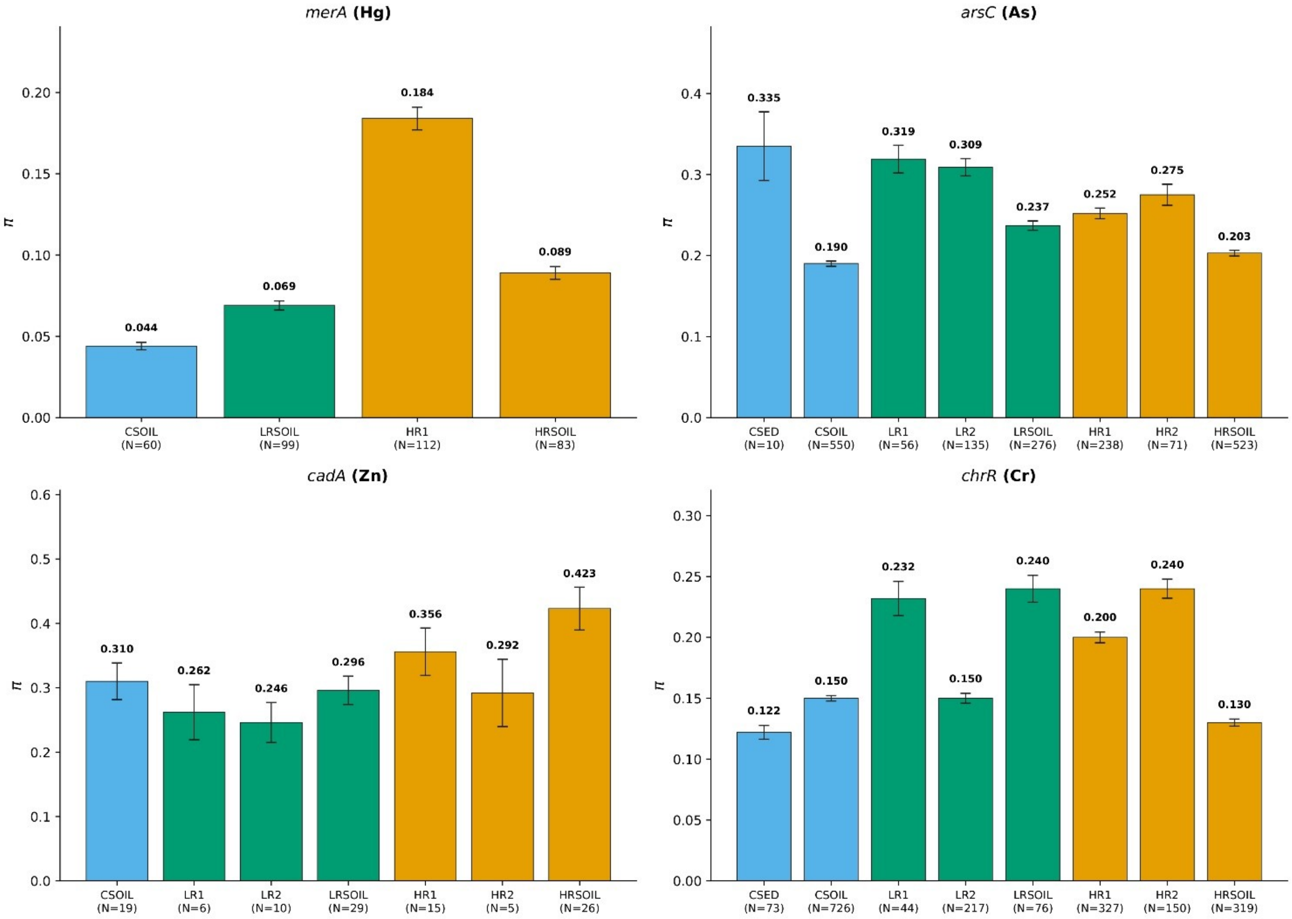


**Supplementary Figure S1.** Nucleotide diversity ($\pi$) per individual sampling site and gene, with sample size N indicated below each bar. Error bars: 95% bootstrap CI. Sites are colored by hydrological regime: Control (blue), Low (green), High (orange).

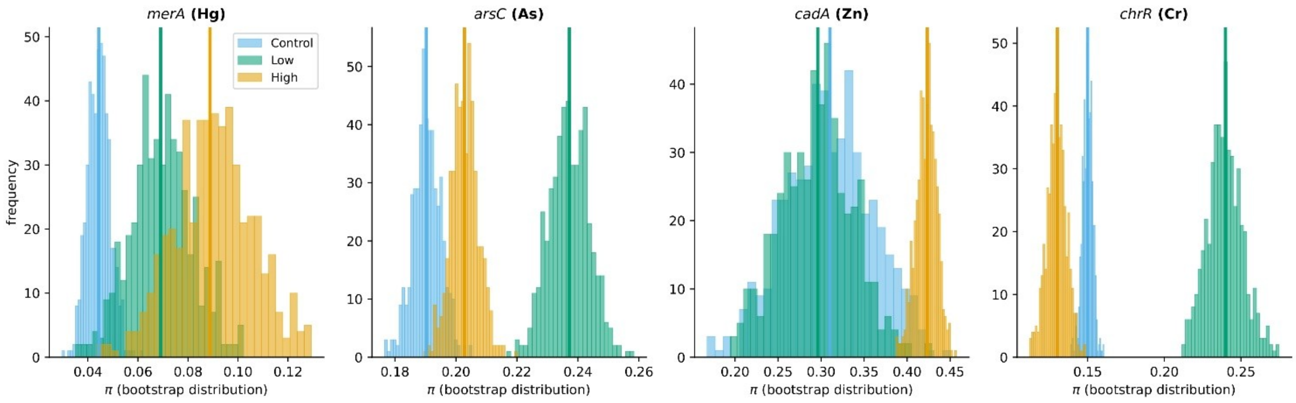


**Supplementary Figure S2.** Bootstrap distributions of nucleotide diversity ($\pi$) per gene and regime, illustrating the precision of point estimates. Vertical lines: point estimates. Distributions are based on 500 bootstrap resamples.

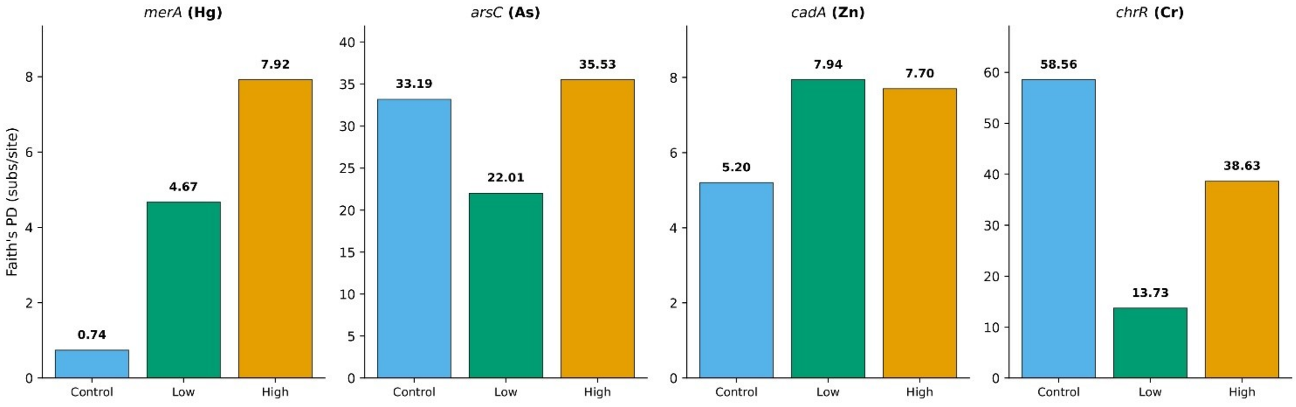


**Supplementary Figure S3.** Faith's phylogenetic diversity (PD) per gene and regime, computed from soil-only IQ-TREE phylogenies. PD = total branch length of the sampled subtree (substitutions per site).

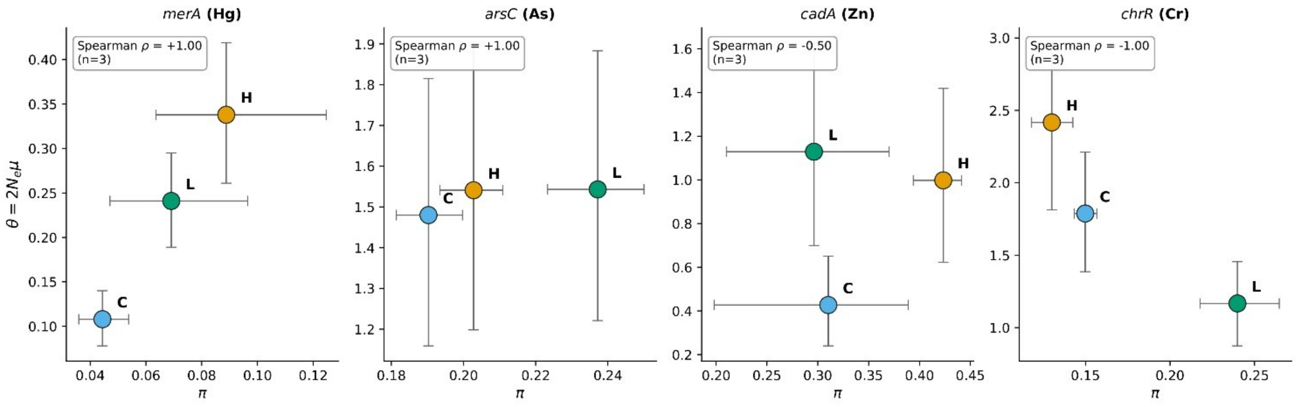


**Supplementary Figure S4.** Relationship between gene-pool diversity index ($\theta = 2Ne\mu$) from BEAST2 and observed nucleotide diversity ($\pi$) per gene and regime. Each point represents one regime. Horizontal error bars: $\pi$ bootstrap 95% CI; vertical error bars: θ 95% HPD.

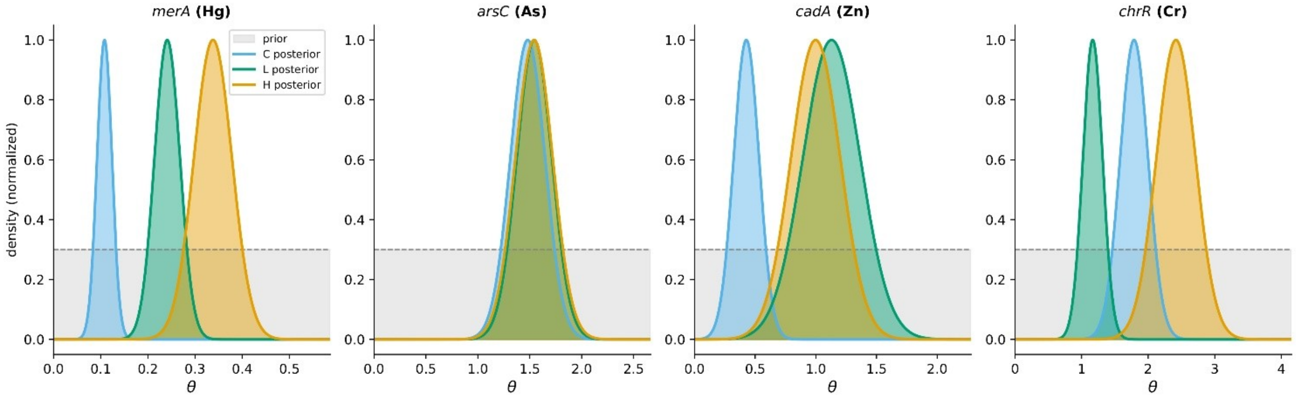


**Supplementary Figure S5.** Prior versus posterior distributions of θ for each gene and regime, illustrating that the data are informative beyond the prior (Exp (1.0)). 95% HPD width reductions of 68 to 98% relative to the prior.

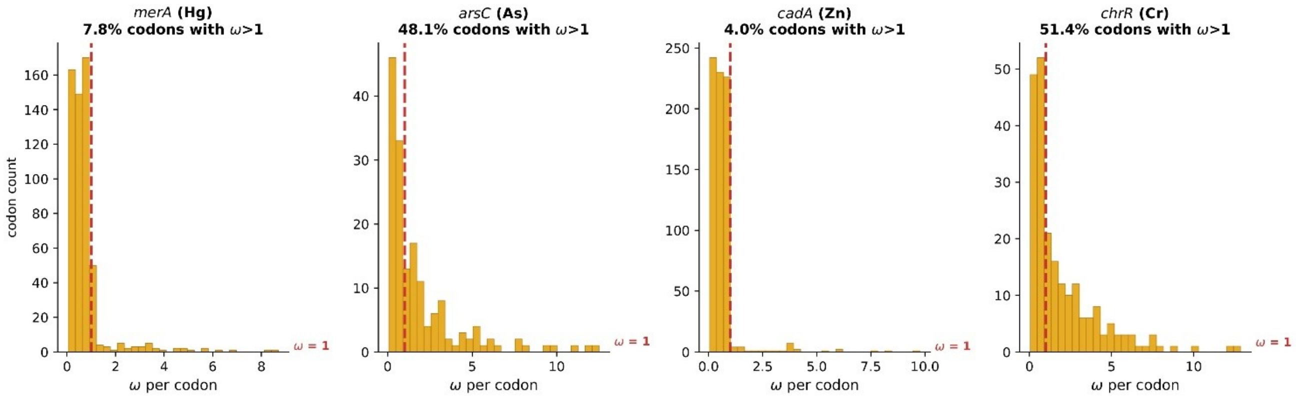


**Supplementary Figure S6.** Distribution of per-codon $\omega = \beta^+/\alpha$ values from MEME for each gene. Red dashed line: $\omega = 1$ (neutral expectation). Percentage of codons with $\omega > 1$ indicated for each gene.
